# Endothelial cells form transient Notch-dependent NO-containing cystic structures during zebrafish cerebrovascular development

**DOI:** 10.1101/416206

**Authors:** E. Kugler, K. Chhabria, S. Daetwyler, J. Huisken, K. Plant, A.M. Savage, R.N. Wilkinson, P.A. Armitage, T.J.A. Chico

**Author notes:** Joint Corresponding Authors Email for correspondence or. Joint Senior Authors (P.A. Armitage, T.J.A. Chico). Authors have declared that no conflict of interest exists.

## Abstract

Endothelial cell behaviour during blood vessel formation is highly complex and dynamic. Transgenic zebrafish have provided many new insights into these processes, due to their ability to provide detailed *in vivo* imaging.

We here report a previously undescribed endothelial cell behaviour during zebrafish embryonic development. Endothelial cells of the cerebral vessels of 3-5d post fertilisation embryos extruded large membranous spherical structures. These were only found on the cerebral vessels, and did not detach from the parent vessel, instead regressing back into the endothelial cell. These structures did not communicate with the vessel lumen, exhibited periodic oscillations in size and shape, and were enriched with filamentous actin at their neck. Due to their unknown nature and spherical appearance we termed these structures *kugeln* (German for sphere).

Pharmacological inhibition of vascular endothelial growth factor (VEGF) signalling significantly increased *kugel* number while Notch inhibition significantly reduced both *kugel* number and diameter. *Kugeln* contain little cytoplasm, but are highly positive for nitric oxide (NO) reactivity, suggesting they represent a novel NO containing organelle specific to the cerebral vessels.

## Introduction

Formation of mature blood vessels requires a wide range of processes, including endothelial proliferation, migration, anastomosis, lumen formation, remodelling, and pruning, alongside recruitment of non-EC types such as vascular smooth muscle cells (Potente and Mäkinen, 2017). Many of these processes have been studied *in vitro*, but only examination of vascular development *in vivo* allows observation of endothelial behaviour within its natural multicellular and complex milieu.

Since visualising embryonic vascular development in mammals is technically challenging the zebrafish has become a widely applied model of vertebrate vascular development. Zebrafish eggs are fertilised externally to the mother, while their translucency enables detailed observation of cellular behaviour during development without instrumentation or *post mortem* histological examination (Gut et al., 2017). An array of transgenic reporter lines that drive fluorescent gene expression in vascular cells, coupled with state-of-the-art imaging techniques such as lightsheet microscopy, enables detailed cellular and subcellular imaging for hours or days (Huisken et al., 2004). This ability to observe vascular development in more detail and for longer periods of time makes it possible to gain new insights into blood vessel formation.

While observing zebrafish cerebrovascular development in transgenic reporter lines by light sheet fluorescence microscopy, we observed a previously undescribed endothelial behaviour restricted to the cerebral vessels which we here describe. Although more work is required to understand the nature of this behaviour, our findings underline the power of unbiased observation of cellular behaviour to reveal novel insights into cell biology.

## Material and Methods

### Zebrafish strains, handling and husbandry

Experiments conformed to UK Home Office regulations and were performed under Home Office Project Licence 70/8588 held by TJAC. Maintenance of adult zebrafish in the Bateson Centre Zebrafish Aquarium of the University of Sheffield and the fish facility at the Max Planck Institute of Molecular Cell Biology and Genetics in Dresden was conducted according to previously described husbandry standard protocols at 28°C with a 14:10 hours light:dark cycle (Westerfield, 1993). Embryos, obtained from controlled pair- or group-mating, were retained in E3 buffer (5mM NaCl, 0.17mM KCl, 0.33mM CaCl_2_, 0.33mM MgSO_4_) with methylene blue.

The following zebrafish lines were used; *Tg(kdrl:HRAS-mCherry)*^*s*916^ labels EC membrane (Chi et al., 2008) (AB background (Streisinger et al., 1981); *Casper* background (White et al., 2008)), *Tg(fli1a:eGFP)*^*y*1^ labels EC cytoplasm (Lawson and Weinstein, 2002), *Tg(flk1:nls-eGFP)*^*zf*109^ labels EC nuclei (Blum et al., 2008), *Tg(fli1a:Lifeact-mClover)*^*sh*467^ labels endothelial filamentous actin (manuscript under review and uploaded to editorial manager), and *Tg(nbt:GCaMP3)* labels neurons (Bergmann et al., 2018).

### Light sheet microscopy of developing zebrafish embryos

Datasets in Sheffield were obtained using a Zeiss Z.1 light sheet microscope with a water-dipping detection-objective (Plan-Apochromat 20×/1.0 Corr nd=1.38); and a scientific complementary metal-oxide semiconductor (sCMOS) detection unit. Data were acquired with activated pivot scan, dual-sided illumination and online fusion; properties of acquired data as follows: 16bit image depth, 1920×1920px image size and minimum z-stack interval (approx. 0.33×0.33×0.5μm). Multi-colour images were acquired in sequential mode. Anaesthetized samples were embedded in 1% or 2% LM-agarose with 0.01% Tricaine in E3 (Sigma). The image acquisition chamber was filled with E3 plus Tricaine, at 28°C.

Light sheet datasets in Dresden were obtained with a home-built multidirectional SPIM (mSPIM) setup (Huisken and Stainier, 2007). The whole head (dorsal) was imaged every 2.5 min over 2 days with dual illumination and 3 μm z-spacing. The mSPIM setup was equipped with a Coherent Sapphire 561nm laser, two Zeiss 10×/0.2 illumination objectives, an UMPlanFL N Olympus 20×/0.5 NA detection objective and an Andor iXon 885 EM-CCD camera. To cover the cerebrovascular region, several regions were imaged and later stitched using custom image processing plugins in Fiji (Schindelin et al., 2012) based on the stitching tool from Stefan Preibisch (Preibisch et al., 2009). Samples were mounted in fluorinated propylene ethylene (FEP) tubes according to established mounting protocols (Kaufmann et al., 2012). The FEP tubes were coated with 3% methyl cellulose and filled with 0.1% low-melting agarose containing 200 mg/l Tricaine to immobilize the zebrafish embryos during time-lapse imaging.

### Microangiography

Vessels perfusion was visualized as previously described (Schmitt et al., 2012; Weinstein et al., 1995), using 20μg dextran tetramethylrhodamine (2,000,000 molecular weight, ThermoFisher) at a concentration of 10mg/μL (2nl injection volume).

### Cessation of blood flow

This was achieved by mechanical opening of heart cavity with forceps. Blood flow cessation was controlled by visual assessment of lack of heart contraction and blood flow.

### Chemical treatments

VEGF signalling was inhibited using the VEGF receptor inhibitor AV951(Nakamura et al., 2006) at 250nM for 2h from 96-98hpf (Selleckchem; S1207; Tivozanib - AVEO pharmaceuticals). Notch signalling was inhibited using 50μM DAPT (Sigma-Aldrich; D4952) for 12h from 84-98hpf (Geling et al., 2002). DMSO was used as control at the same concentration and duration for both drug treatments.

### Vital dye staining

*In vivo* visualization of nitric oxide was performed via application of 2.5μM DAF-FM-DA (Molecular Probes; D23844) (Kojima et al., 1998) for 6h in 4dpf embryos (96-102hpf). DMSO control was performed at the same concentration and duration. To visualize acidic cellular compartments LysoTracker Green (Molecular Probes; L7526 DND-26) was applied for 5h (96-101hpf) at a concentration of 8.33μM in E3 (He and Klionsky, 2010).

### Image analysis and representation

Images were analysed using open-source software Fiji (Schindelin et al., 2012). Kymographs were produced via stack reslicing to study *kugel* diameter changes over time. To visualize data, maximum intensity projections (MIP) were generated and displayed using either gray (single channel) or red/green (multi-channel) colour representations. Intensity inversion was applied, as appropriate, to give the clearest rendering of relevant structures. Time-lapse data were displayed using MIPs.

### Statistical analysis

Normality of data was tested using D’Agostino-Pearson omnibus test. Statistical analysis of normally distributed data was performed using a One-way ANOVA to compare multiple groups or Students t-test to compare two groups. Non-normally distributed data were analysed with a Kruskal-Wallis test to compare multiple groups, or Mann-Whitney test to compare two groups. Analysis was performed in GraphPad Prism Version 7 (GraphPad Software, La Jolla California USA). P values are indicated as follows: p<0.05 *, p<0.01 **, p<0.001 ***, p<0.0001 ****. Data represents mean and standard deviation, if not otherwise stated. Image representation was performed using Inkscape (https://www.inkscape.org).

## Results

We used light sheet fluorescence microscopy to observe formation of the embryonic cerebral vasculature in *Tg(kdrl:HRAS-mCherry)*^*s*916^ zebrafish expressing an endothelial membrane-tagged reporter protein (Chi et al., 2008). While analysing these datasets we frequently observed rounded protrusions arising from the ECs of the cerebral vessels (**Figure 1A**). Although their appearance on single z-slices or MIPs was similar to cross-sections of lumenized vessels, detailed imaging and three-dimensional reconstruction revealed these structures to be blind-ended and spherical (**Figure 1B, Supplemental Video 1**). Such structures were observed in datasets generated in both Sheffield and Dresden using unrelated zebrafish colonies. The mean±SEM diameter of these structures at 3dpf was 10.1±0.5μm (**Figure 1C**), which is far larger than microvesicles or other previously described membrane derived structures, and similar to, or exceeding, the luminal diameter of the parent vessel from which they arose. Due to their large size, spherical shape, and the absence of an existing term to describe such structures we named them *kugeln* (singular *kugel*).

**Figure 1.**
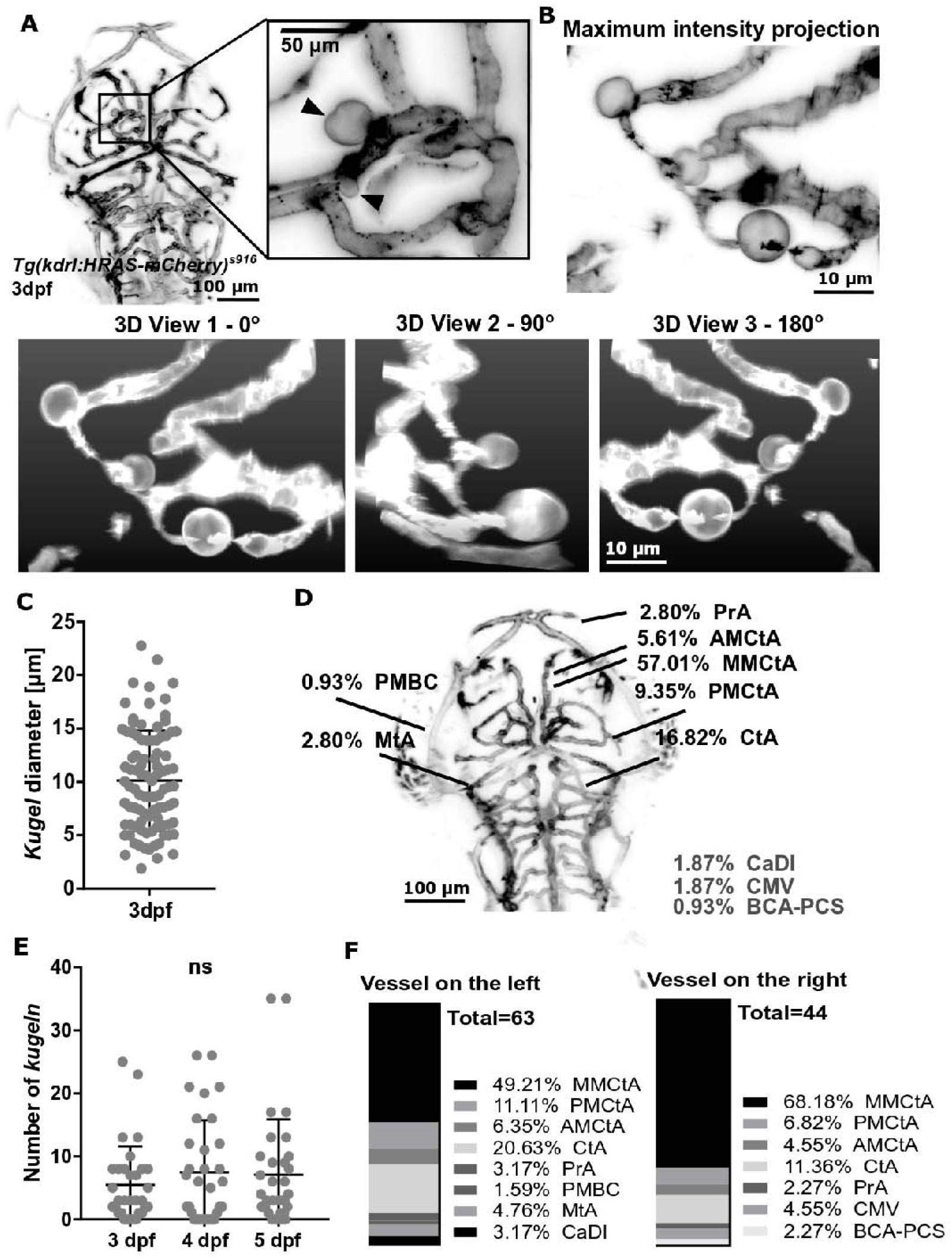
ECs of the cerebral vessels of developing zebrafish often display large spherical protrusions which we term “*kugeln”*. **A** MIP of cerebral vessels of 3dpf *Tg(kdrl:HRAS-mCherry)*^*s*916^ embryos (inverted). Higher magnification panel showed two endothelial membrane protrusions arising in the middle mesencephalic central artery (MMCtA) (arrowheads). **B** 3D reconstruction showed *kugeln* were spherical protrusions from the parent MMCtA. **C** The diameter of *kugeln* at 3dpf (n=93 from 32 embryos; 3 experimental repeats). **D** Site and location of *kugeln* (n=107 from 34 embryos at 4dpf; 3 experimental repeats). *AMCtA* – anterior mesencephalic central artery, *BCA* – basal communicating artery, *CaDI* – caudal division of internal carotid artery, *CMV* – communicating vessel, *CtA* – central arteries, *dpf* – days post fertilization, *L* – left, *MMCtA* – middle mesencephalic central artery, *MtA* – metencephalic artery, *PCS* – posterior communicating segment, *PMBC* – posterior midbrain channel, *PMCtA* – posterior mesencephalic central artery, *PrA* – prosencephalic artery, *R* - right. **E** Number of *kugeln* per embryo between 3-5dpf (3dpf: 93 *kugeln,* 32 embryos; 4dpf: 81 *kugeln,* 32 embryos, 5dpf: 73 *kugeln,* 32 embryos; 3 experimental repeats). **F** Location of *kugeln* by vessel and laterality (4dpf 107 *kugeln*, 34 embryos; 3 experimental repeats).

We found over 90% of *kugeln* were located on the central vessels of the cerebral vasculature (**Figure 1D**). From 3-5dpf, we observed *kugeln* on cerebral vessels of most *Tg(kdrl:HRAS-mCherry)*^*s*916^ embryos as follows: 3dpf in 33/39 fish, 4dpf 26/37 fish and 5dpf 29/37 fish. The mean number of *kugeln* per embryo did not differ between 3-5dpf (**Figure 1E**; Kruskal Wallis p 0.8571). Although individual *kugeln* were often unilateral (present on left-side but not right-sided vessels, or vice versa) overall left/right distribution was not significantly different (**Figure 1F**; Wilcoxon test p 0.0605).

We next acquired time-lapses to observe development and behaviour of *kugeln*. The resulting time-lapse data revealed the extrusion and retraction of *kugeln* (**Figure 2A**). Though we expected *kugeln* would form separate vessels or interact with other ECs to anastomose, *kugeln* always either regressed back into the parent vessel or persisted to the end of the dataset without separation or anastomosis. The lifetime of *kugeln* was variable. While some persisted for minutes, others remained stable for hours (**Figure 2B**). Further examination showed that *kugeln* display dynamic alteration in shape and size, including shape changes, enlargement and retraction (**Figure 2C and Supplemental Video 2**). Kymographs were used to visualize diameter changes of *kugeln* and cerebral vessels over a 20min time period. This revealed an oscillatory behaviour of the diameter of *kugeln* with a periodicity of minutes (**Figure 2D**). No such alteration was present in the diameter of the cerebral vessel, suggesting that changes of *kugel* size or shape were not due to changes in blood pressure which has a periodicity of milliseconds.

**Figure 2.**
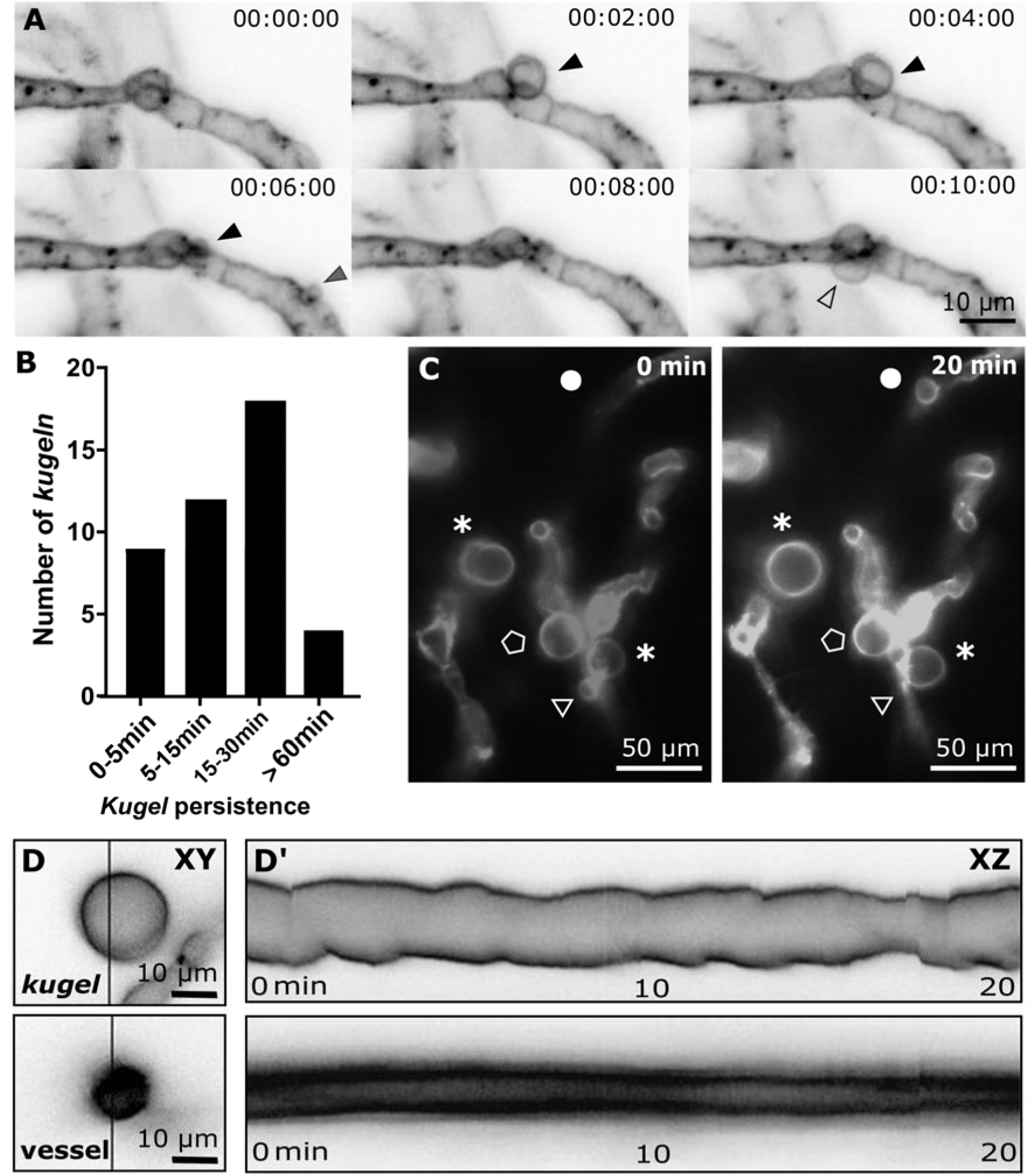
Endothelial *kugeln* are transient and dynamically alter shape and size. **A** MIPs of a timeseries light sheet acquisition at 2min intervals (inverted). Three *kugeln* indicated by black, grey and unfilled arrowheads. **B** *Kugeln* persisted for variable durations before regression into the parent vessel (4dpf 43 *kugeln*, 9 embyros; 4 experimental repeats). **C** During their lifetime *kugel* behaviours included shape changes (asterisk), retraction (circle), expansion (triangle), while others changed little in the same period (pentagon). **D** Kymographs generated by line-scanning across the diameter of a typical *kugel* or a similarly sized cerebral vessel revealed that kugel diameter oscillated with a periodicity of minutes.

We next investigated the cellular nature and composition of endothelial *kugeln*. We examined transgenic embryos that label endothelial nuclei, and identified no *kugeln* containing a nucleus (**Figure 3A**). This, coupled with no evidence of shedding from the parent cell, suggests that *kugeln* do not represent an atypical form of cell proliferation or apoptosis. When we examined a transgenic labelling filamentous actin *Tg(fli1a:LifeAct-mClover)*^*sh*467^ in ECs, we observed that *kugeln* were enriched for filamentous actin (F-actin) especially at the *kugel* “neck” (**Figure 3B** and **Supplemental Video 3**). This actin enrichment and the dynamic nature of *kugel* development, oscillation and regression suggest cytoskeletal rearrangement may be necessary for their growth and/or maintenance. To characterise the contents of *kugeln*, we examined double transgenics expressing both membrane labelling *Tg(kdrl:HRAS-mCherry)*^*s*916^ (Chi et al., 2008) and cytosolic GFP using *Tg(fli1a:eGFP)*^*y*1^ (Lawson and Weinstein, 2002). We were unable to visualise cytosolic GFP in the majority of *kugeln*; most were not visible using the cytosolic reporter and were only observed using the membrane-tagged reporter (**Figure 3C**). To characterise the tissue surrounding the *kugeln*, we examined double transgenics expressing GFP in neurons of the developing brain (Bergmann et al., 2018), and observed exclusion of neural tissue from *kugeln* (**Figure 3D**).

**Figure 3.**
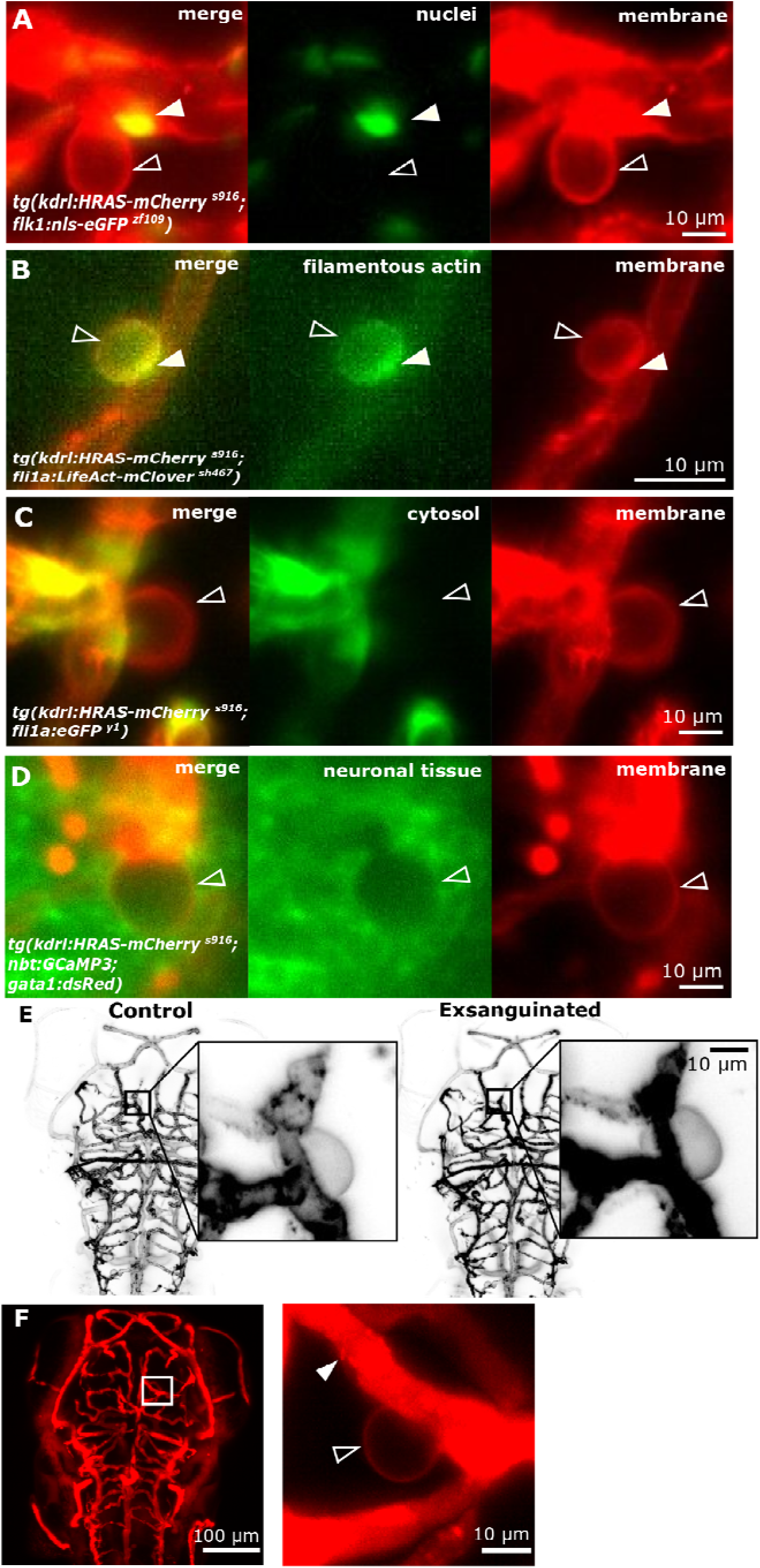
Endothelial *kugeln* are non-nucleated, F-actin-rich and do not communicate with the parent vessel lumen. **A** Double transgenic visualizing endothelial membrane (red) and endothelial nuclei (green). A nucleus (arrowhead) was observed close to but not within the *kugel* (unfilled arrowhead). **B** Double transgenic showing endothelial membrane (red) and endothelial F-actin (green). F-actin localized to the neck of the *kugel* (arrowhead). **C** Double transgenic showing endothelial membrane (red) and endothelial cytoplasm (green); the cytoplasmic reporter was visible in the parent vessel but not in the *kugel* (unfilled arrowhead). **D** Double transgenic showing endothelial membrane (red) and neurons (green). Surrounding neurons were excluded from the volume of the *kugel* (unfilled arrowhead). **E** MIPs of the cerebral vessels of an individual 4 dpf embryo before and after exsanguination showing that this did not alter *kugel* size. **F** Dextran microangiography found dextran filled all lumenized vessels (arrowhead) but did not enter the *kugel* (unfilled arrowhead).

We next examined the relationship between endothelial *kugeln* and the lumen of the parent vessel. We identified *kugeln* in individual animals, which were then exsanguinated by cardiac puncture to abolish blood flow and blood pressure (**Figure 3E**). This had no effect on *kugel* size or shape, suggesting neither blood flow nor pressure is needed to maintain *kugeln* once formed. We next performed microangiography with fluorescent dextran. No entry of dextran into *kugeln* was observed, confirming a lack of connection between the lumen of the parent vessel and the contents of the *kugel* after formation (**Figure 3F**).

VEGF and Notch signalling are central orchestrators of vascular development(Gerhardt et al., 2003; Gore et al., 2012; Siekmann and Lawson, 2007). We therefore examined the effect of pharmacological inhibition of these signalling pathways on *kugel* formation. Treatment with the VEGF inhibitor, AV951, significantly increased the number of *kugeln* per embryo, without affecting *kugel* diameter (**Figures 4A-C**). In contrast, Notch inhibition with DAPT significantly reduced both number of *kugeln* per embryo and *kugel* diameter (**Figures 4D-F**). To better understand the contents of endothelial *kugeln,* we stained embryos with the vital dye DAF-FM, which fluoresces green in contact with nitric oxide (Kojima et al., 1999, 1998). After DAF-FM staining 63% of *kugeln* (37/59 *kugeln*, 12 embryos) were filled with fluorescent dye, suggesting *kugeln* contain significant levels of NO (**Figure 4G**). We performed similar experiments with LysoTracker to visualise acidic cell compartments and found that 23% of *kugeln* (55/240 *kugeln,* 22 embryos) (**Figure 4H**). To understand at which point in the lifetime of a kugel it begins to contain NO we performed time-lapse microscopy of *kugel* formation after DAF-FM staining and found this to be present at commencement or even just prior to *kugel* formation (**Figure 4I**).

**Figure 4.**
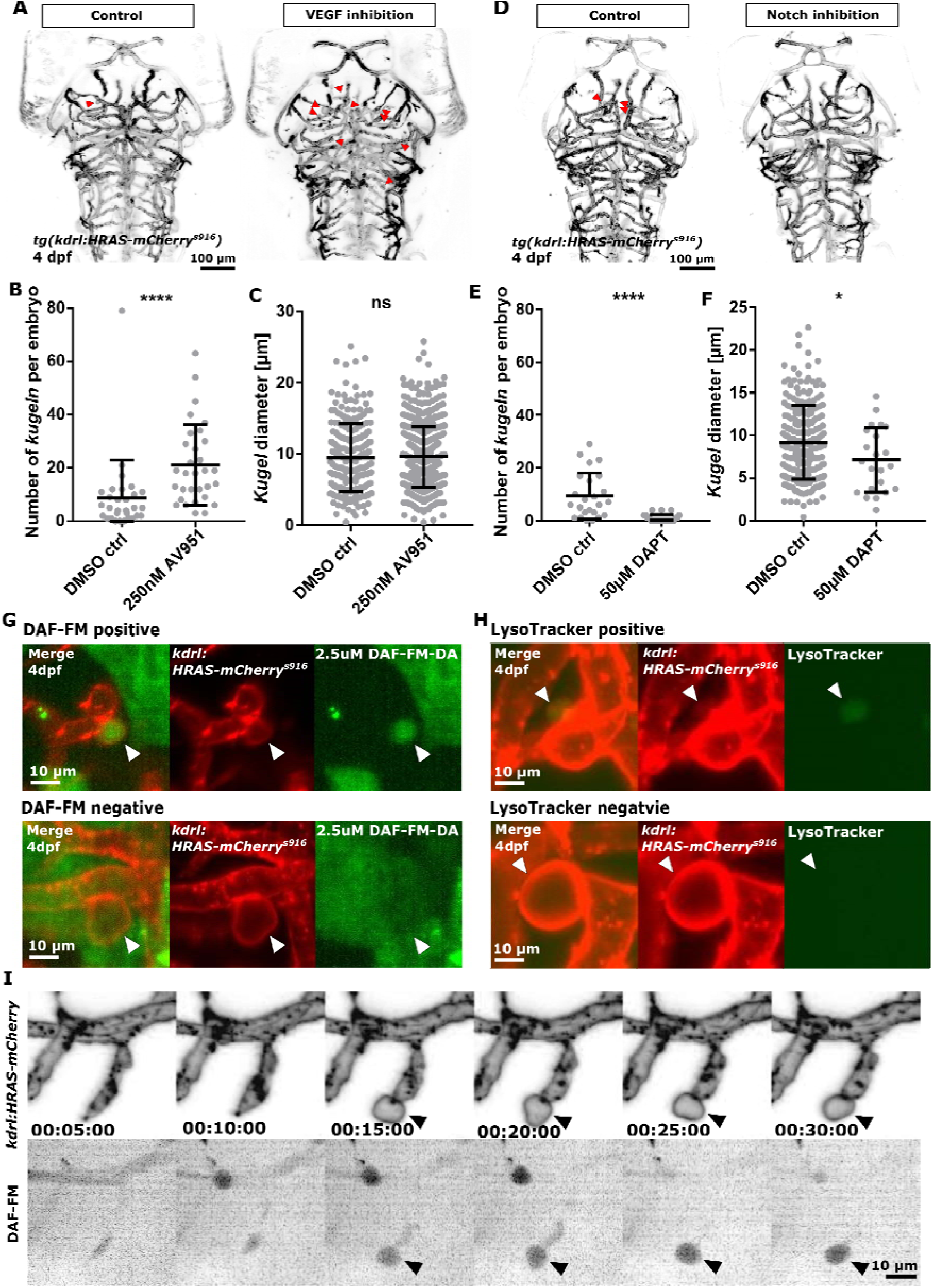
Endothelial *kugeln* contained NO, and *kugeln* number was increased by VEGF inhibition and reduced by Notch inhibition. **A** Representative 4dpf zebrafish embryos incubated with DMSO control or the VEGF inhibitor AV951 at 250nM for 2h from 96-98hpf; *kugeln* indicated by red arrowheads. **B** VEGF inhibition significantly increased *kugel* number (p<0.0001; ctrl n=30 larvae; AV951 n=31 larvae from 4 experimental repeats; Mann-Whitney U test). **C** Mean *kugel* diameter was not statistically significantly different after AV951 exposure (p 0.4034; ctrl n=243 *kugeln* from 30 larvae; AV951 n=652 *kugeln* from 31 larvae; 4 experimental repeats; Mann-Whitney U test). **D** Representative 4dpf zebrafish embryos incubated with DMSO control or Notch inhibitor DAPT at 50μM for 12h from 84-98hpf; *kugeln* indicated by red arrowheads. **E** *Kugel* number was significantly decreased by inhibition of Notch signalling (p<0.0001; n=24 larvae from 3 experimental repeats; Mann-Whitney U test). **F** Mean *kugel* diameter was significantly reduced by DAPT treatment (p 0.0359; ctrl n=225 *kugeln* from 24 larvae; DAPT n=22 from 24 larvae; 3 experimental repeats; Mann-Whitney U test). **G** DAF-FM staining (2.5μM incubation from 96-102hpf) showed that 62.72% of *kugeln* were filled with nitric oxide (n=12 4dpf larvae; 37/59 *kugeln* filled; 2 experimental repeats). Images showed representative “filled” (DAF-FM positive) and “unfilled” (DAF-FM negative) *kugeln*. **H** Application of LysoTracker, a vital dye that stains lysosomes or acidic compartments (8.33μM; 96-101hpf) found that 23% of *kugeln* contained acidic contents. Images showed representative “filled” (Lysotracker positive) and “unfilled” (Lysotracker negative) *kugeln*. (n=22 4dpf larvae; 55/240 *kugeln* filled; 2 experimental repeats). **I** Time-lapse acquisition with DAF-FM revealed that *kugeln* contained NO early in their biogenesis.

## Discussion

Lipid membrane vesicle formation of ECs has been well characterised, comprising three subgroups with different diameters: apoptotic bodies (1-4µm), microvesicles (0.15-1µm) and exosomes (40-150nm) (Colombo et al., 2014). We here describe an as yet unreported form of EC behaviour in zebrafish found only on the cerebral vessels in which EC develop large spherical protrusions that oscillate in shape and regress. *Kugeln*, with mean diameter of 10µm are far larger than these vesicles, and never separate from their parent vessel. Therefore, although the biogenesis of *kugeln* may share common mechanisms with these vesicles, these two cardinal features (size and lack of shedding) support our contention that *kugeln* represent a different phenomenon.

Transgenic zebrafish are an established model of vascular development, having contributed greatly to our understanding of the mechanisms that govern blood vessel formation. Given the large number of published studies using the model, we were surprised to find no prior reports of *kugeln*. This is likely to be due to the fact that *kugeln* were only detected using membrane-tagged reporter proteins, while *kugeln* were not visible using more widely used cytoplasmic reporter lines such *as Tg(fli1a:eGFP)*^y1^ (Lawson and Weinstein, 2002). In addition, most studies of vascular development in zebrafish have examined the trunk vasculature, in which we found no evidence of *kugeln*. Our findings indicate that EC behaviour may differ markedly between vascular territories even at the same developmental stage in the same animal.

VEGF and Notch signalling are key drivers of vascular development, acting in a highly co-ordinated and context-dependent manner to orchestrate endothelial behaviour. Notch signalling generally reduces sensitivity to VEGF signalling by mechanisms including reduction of VEGF receptor expression (Thomas et al., 2013). Neither VEGF nor Notch have previously been implicated in vesicle biogenesis of any cell type. Our findings suggest that Notch signalling in developing cranial endothelial cells may function as part of the regulation of *kugel* formation. However, further understanding of how downstream signalling and feedback mechanisms influence *kugel* formation is required.

Our finding that the majority of *kugeln* contained high levels of NO was unexpected as NO is generally considered to passively diffuse through membranes from cells producing it (Thomas, 2015). We could not find any previous description of NO-containing organelles. The link between VEGF, Notch, and *kugeln* is unclear; VEGF induces NO release from ECs (Ziche et al., 1997) but appears to inhibit *kugeln* formation. Zebrafish do not possess a clear orthologue of endothelial nitric oxide synthase (eNOS). However, zebrafish do possess orthologues for inducible and neuronal NOS and it may be these isoforms that generated the NO we detected (Lepiller et al., 2007).

Kugeln do not represent a form of aneurysm, since they do not directly connected to the vascular lumen, and their maintenance does not require blood flow or pressure. This observation, and the fact that *in vivo* imaging revealed F-actin was highly enriched at the neck of *kugeln*, suggests their formation is due to active cytoskeletal rearrangements rather than passive “inflation”.

Many important questions regarding *kugeln* remain, particularly to determine their function and mechanism of biogenesis, the reason for their restriction to cerebral vessels, and whether they exist in other species. Given the conservation of endothelial behaviour between zebrafish and mammals, and the fact that only *in vivo* imaging of membrane-tagged reporters would be able to distinguish *kugeln* from vessel lumens, we consider it highly likely that we have uncovered a behaviour that is not restricted to zebrafish but which may provide further insight into the remarkable range of EC behaviours during vascular development.

## Acknowledgements

We thank the Bateson Centre aquarium facility for excellent zebrafish husbandry. This work was supported by a University of Sheffield, Department of Infection, Immunity and Cardiovascular Disease, Node Studentship. The Zeiss Z1 LSFM was funded via British Heart Foundation Infrastructure Award IG/15/1/31328 awarded to CT and RW.

## Author Contribution

Funding Acquisition, PA, TC, RW, JH; Investigation, Validation and Data Curation, EK, SD, KP, and KC; Formal Visualization and Analysis, EK; Resources, AP, TC, AS, RW, and JH; Project Administration, EK, KP, PA, and TC; Writing – Original Draft, EK, and TC; Writing – Review and Editing, all authors contributed equally;

## Supplemental Video Legends

Supplemental Video 1. 3-dimensional reconstruction of cerebral vessels of *Tg(kdrl:HRAS-mCherry)*^*s*916^ transgenic zebrafish shown in Figure 1B.

Supplemental Video 2. Timelapse lightsheet microscopy of *kugeln* arising from cerebral vessels of *Tg(kdrl:HRAS-mCherry)*^*s*916^; *Tg(flk1:nls-eGFP)*^*zf*109^ transgenic zebrafish.

Supplemental Video 3. Timelapse lightsheet microscopy of *kugeln* arising from cerebral vessels of *Tg(kdrl:HRAS-mCherry)*^*s*916^; *Tg(fli1a:Lifeact-mClover)*^*sh*467^ transgenic zebrafish.

